## Supplementary Figures for "Shifting the PPARγ conformational ensemble towards a transcriptionally repressive state improves covalent inhibitor efficacy"

File contains:

- Figure 6—figure supplement 1-4
- Figure 7—figure supplement 1

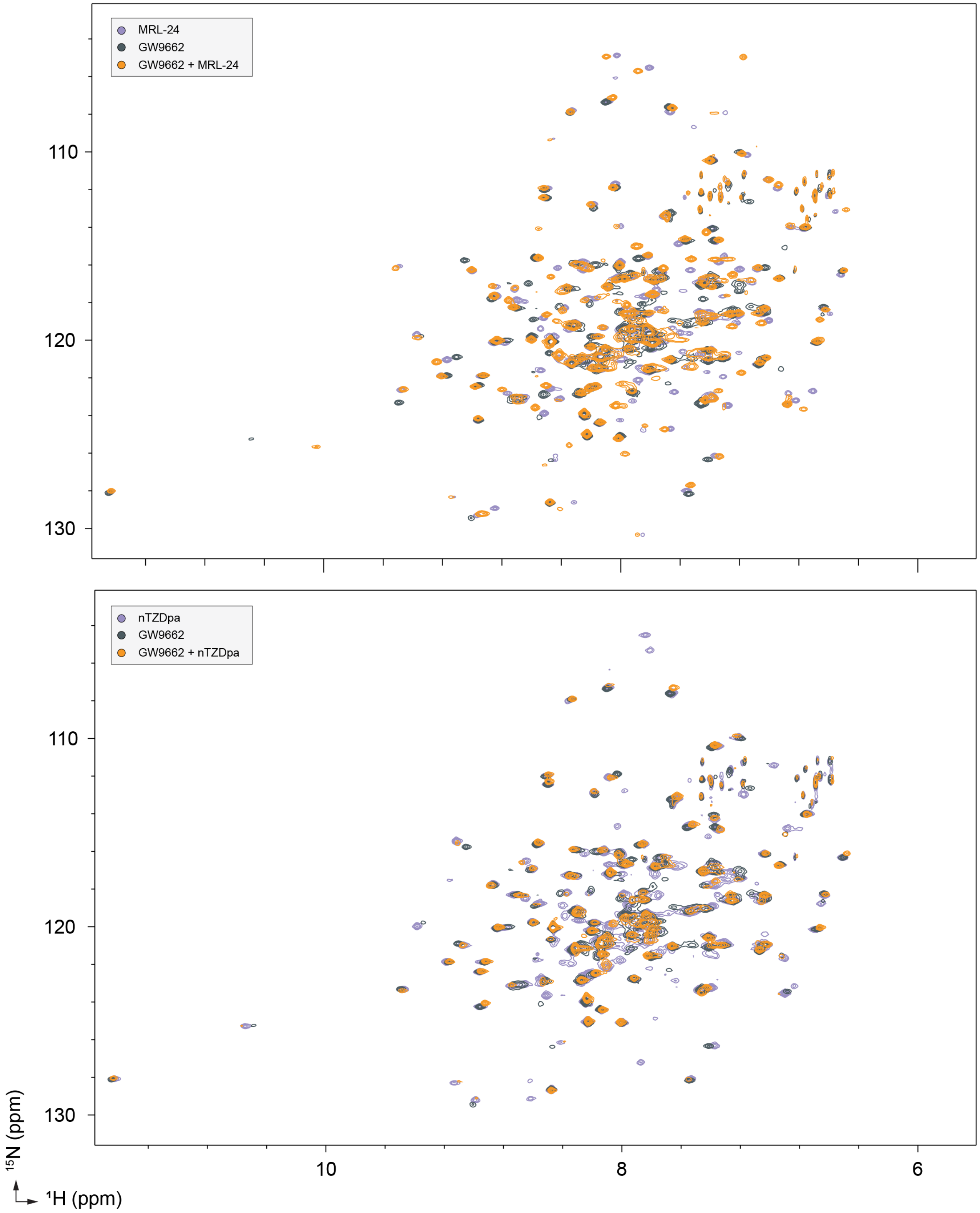


**Figure 6—figure supplement 1.** 2D [$^{1}$H,$^{15}$N]-TROSY-HSQC NMR data of $^{15}$N-labeled PPARγ LBD bound individually or cobound to MRL-24 (top), nTZDpa (bottom), and SR16832 as indicated.

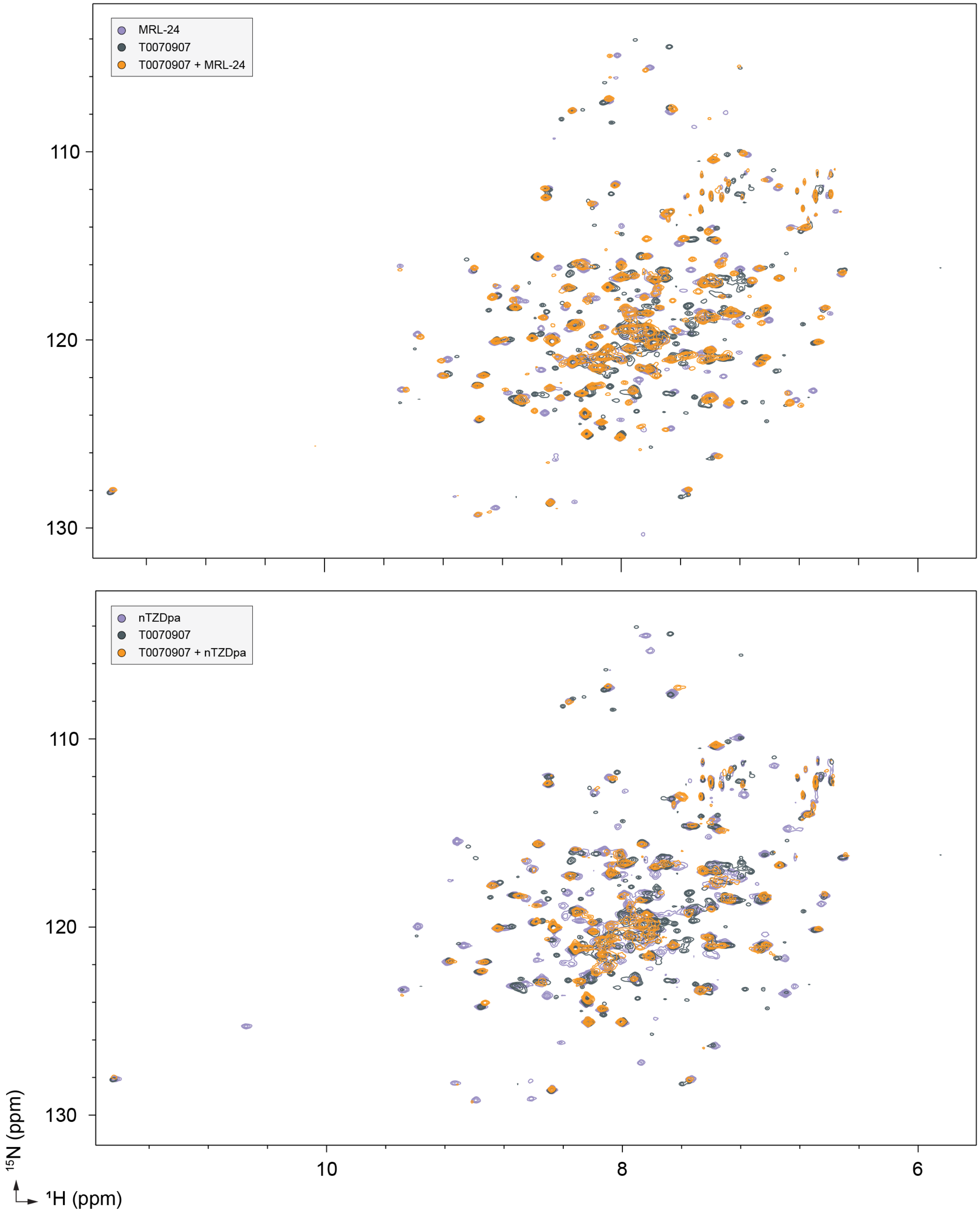


**Figure 6—figure supplement 2.** 2D [$^{1}$H,$^{15}$N]-TROSY-HSQC NMR data of $^{15}$N-labeled PPARγ LBD bound individually or cobound to MRL-24 (top), nTZDpa (bottom), and T0070907 as indicated.

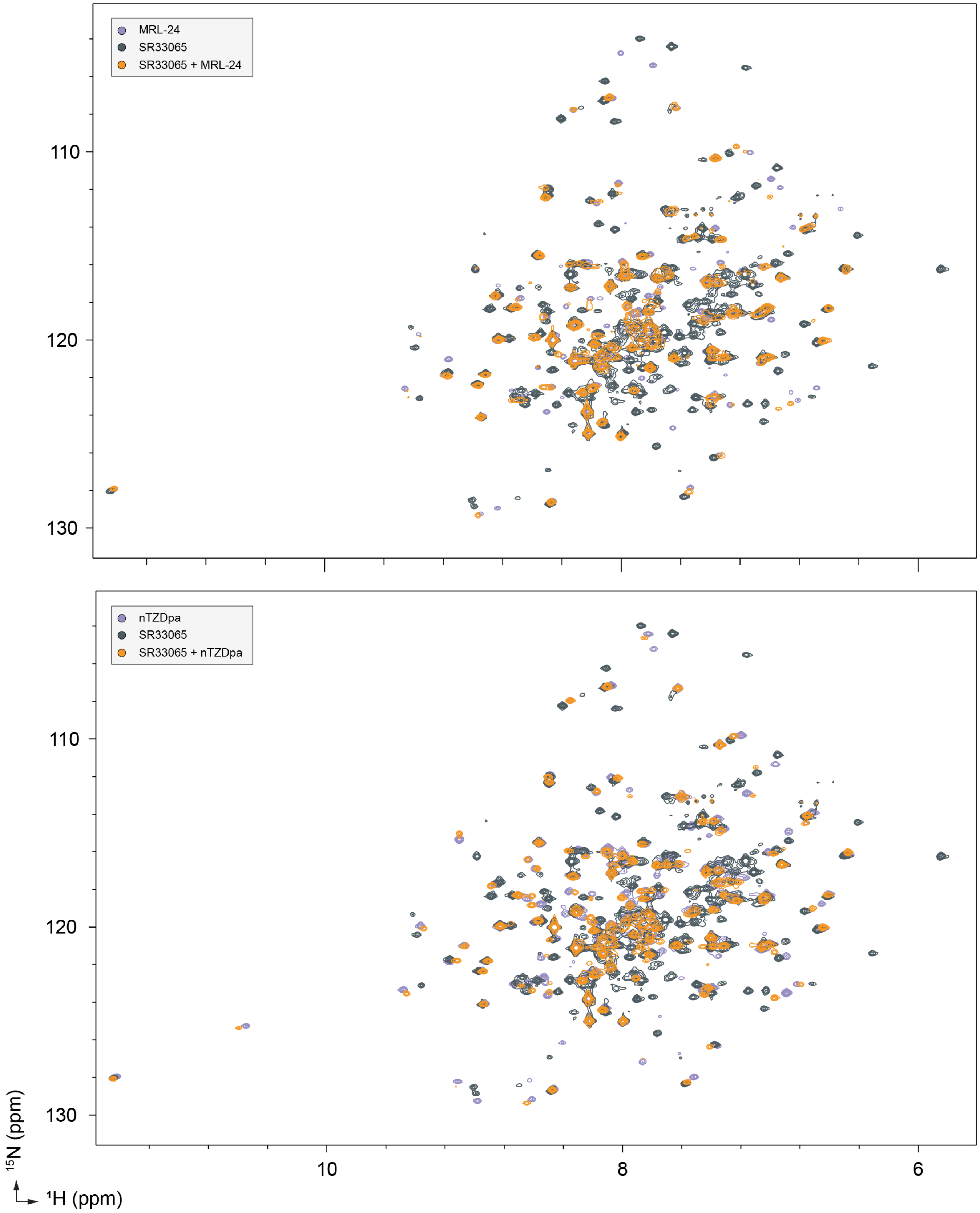


**Figure 6—figure supplement 3.** 2D [$^1$H,$^{15}$N]-TROSY-HSQC NMR data of $^{15}$N-labeled PPARγ LBD bound individually or cobound to MRL-24 (top), nTZDpa (bottom), and SR33065 as indicated.

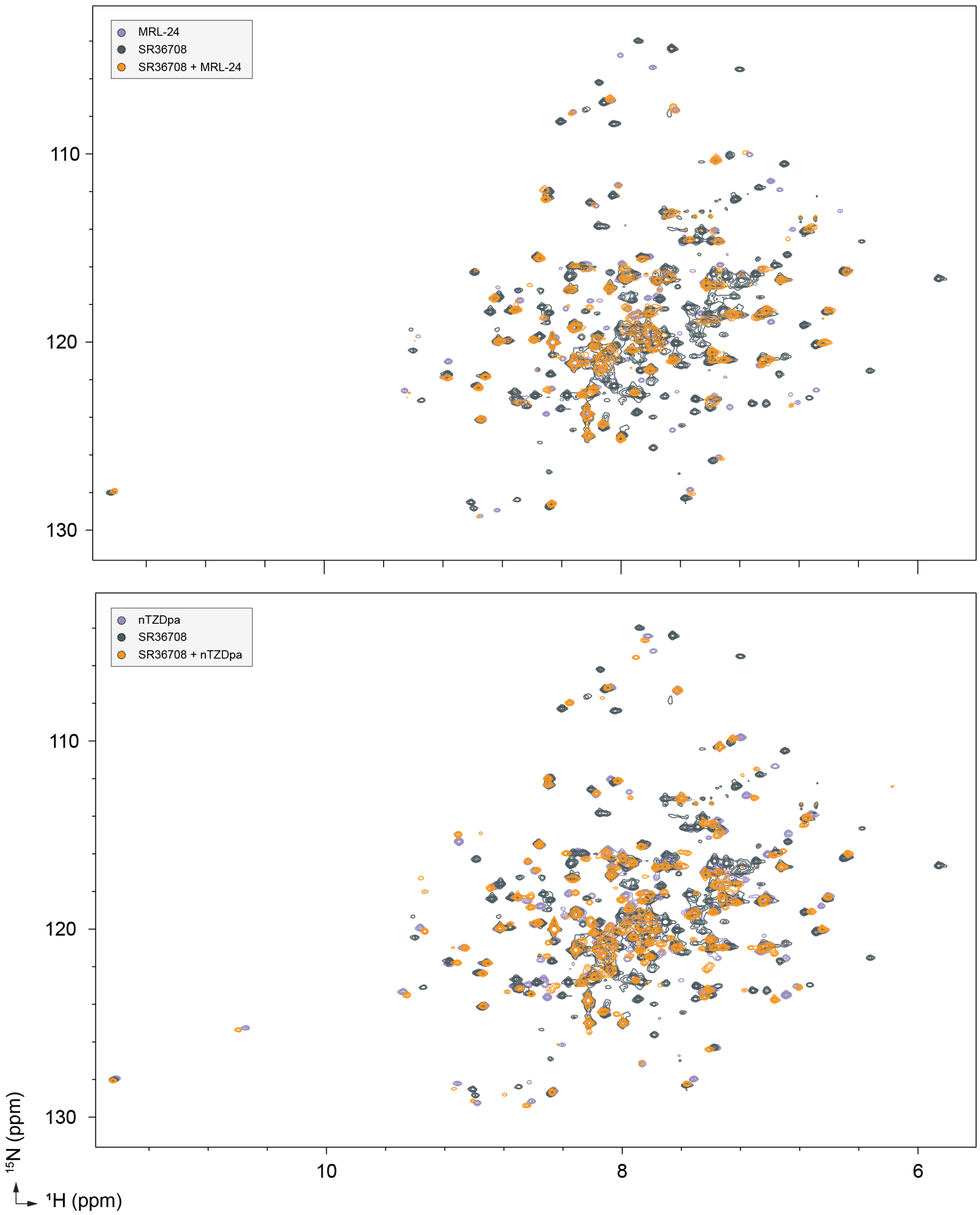


**Figure 6—figure supplement 4.** 2D [$^{1}$H,$^{15}$N]-TROSY-HSQC NMR data of $^{15}$N-labeled PPARγ LBD bound individually or cobound to MRL-24 (top), nTZDpa (bottom), and SR36708 as indicated.

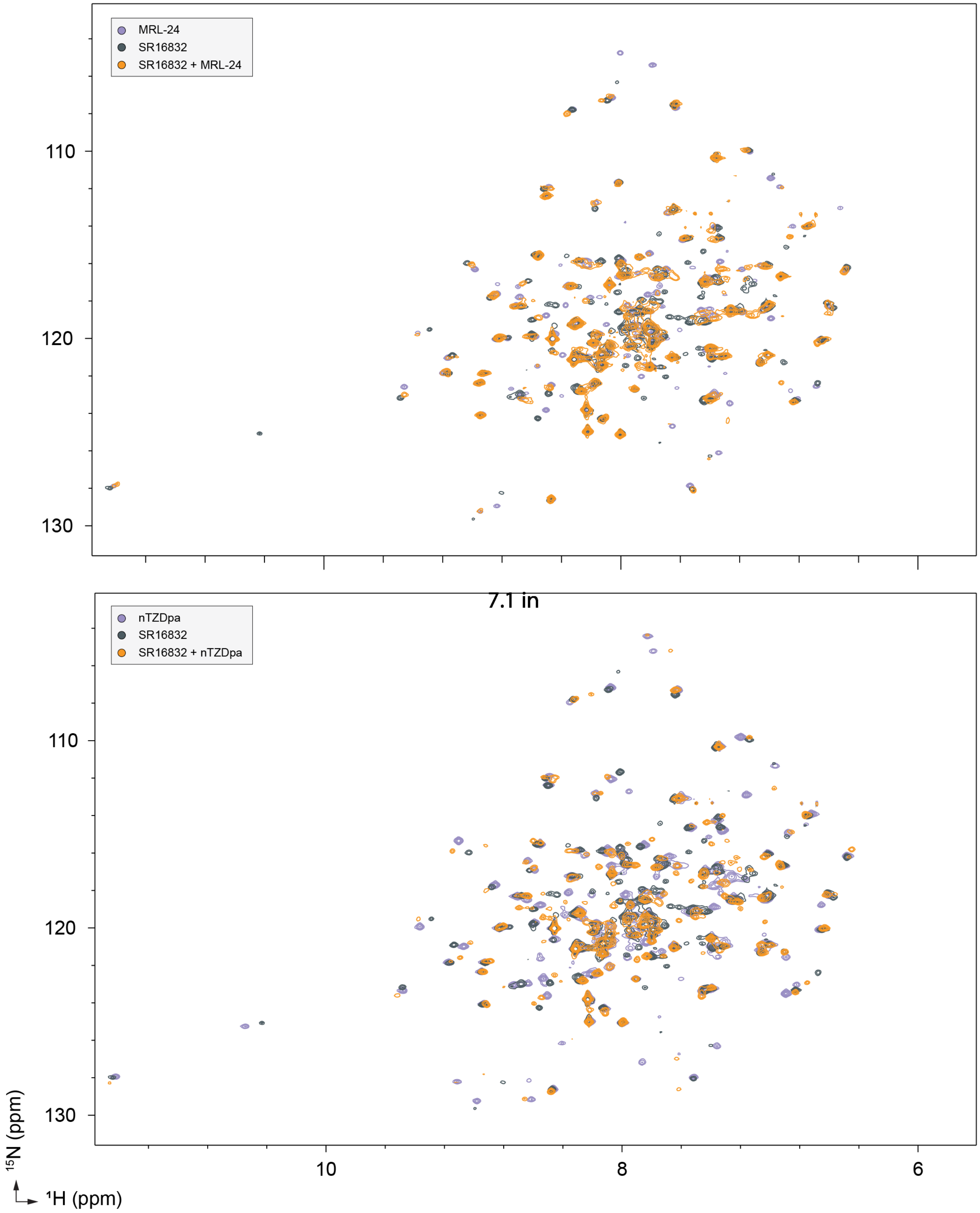


**Figure 7—figure supplement 1.** 2D [$^{1}$H,$^{15}$N]-TROSY-HSQC NMR data of $^{15}$N-labeled PPARγ LBD bound individually or cobound to MRL-24 (top), nTZDpa (bottom), and SR16832 as indicated.
